# Rice leaves undergo a rapid metabolic reconfiguration during a specific stage of primordium development

**DOI:** 10.1101/2021.12.01.470706

**Authors:** Naomi Cox, Heather J. Walker, James K. Pitman, W. Paul Quick, Lisa M. Smith, Andrew J. Fleming

## Abstract

Leaf development is crucial to establish the photosynthetic competency of plants. It is a process that requires coordinated changes in cell number and differentiation, transcriptomes, metabolomes and physiology. However, despite the importance of leaf formation for our major crops, early developmental processes for rice have not been comprehensively described. Here we detail the temporal developmental trajectory of early rice leaf development and connect morphological changes to metabolism. In particular, a developmental index based on the patterning of epidermal differentiation visualised by electron microscopy enabled high resolution staging of early growth for single primordium metabolite profiling. These data demonstrate that a switch in the constellation of tricarboxylic acid (TCA) cycle metabolites defines a narrow window towards the end of the P3 stage of leaf development. Taken in the context of other data in the literature, our results substantiate that this phase of rice leaf growth, equivalent to a change of primordium length from around 5 to 7.5 mm, defines a major shift in rice leaf determination towards a photosynthetically defined structure. We speculate that efforts to engineer rice leaf structure should focus on the developmental window prior to these determining events.

**Highlight:** Rice leaves undergo a shift in fundamental metabolism during a very early and narrow developmental window which co-incides with them acquiring the ability to capture light for photosynthesis

## INTRODUCTION

Leaves develop from a shoot apical meristem which is non-photosynthetic. Therefore, at some stage a switch to photosynthetic competence must occur so that the mature leaf can provide the carbohydrates required for further plant growth and development (Fleming, 2006). To achieve this, co-ordinated processes of cell division, elongation and differentiation occur as the leaf passes through a series of developmental stages from an initial primordium. These leaf stages can be defined by plastochron number, (Pi) (P1, P2, P3…), which describes the developmental age of the leaf independent of time, with P1 being the stage subsequent to initiation up to the point at which the next primordium is generated by the meristem (Erickson and Michelini, 1957). At this point the P1 stage primordium is defined as entering the P2 stage of development. As new primordia are formed at the meristem, the first primordium passes through developmental stages P3, P4 and so on. Thus each leaf can be defined both by the order of initiation (1^st^, 2^nd^, 3^rd^, …) and by developmental stage (P1, P2, P3, …), providing a robust platform for developmental analysis. Although there have been studies providing detailed accounts of the earliest stages of leaf development in a number of plant species, e.g., Arabidopsis (Kalve *et al*., 2014), maize (Wang *et al*., 2013) and *Cardamine hirsuta* (Hay and Tsiantis, 2016), detailed information on the earliest stages of rice leaf development remains fragmented or superficial. For example, one of the most comprehensive studies (Itoh *et al*., 2005) provides an excellent description of the entire structural process of leaf development but, of necessity, provides more limited detail on the very earliest stages and, moreover, lacks information on the accompanying changes at the level of transcriptome, metabolome and physiology. A recent study by Miya and colleagues (2021) analysed changes in the transcriptome over rice leaf development from very early leaf development through to mature leaf blades using microarrays (Miya *et al*., 2021), building on a previous RNA-seq analysis that specifically focused on the P3 to P4 transition to photosynthetic competence in leaf 5 (van Campen *et al*., 2016). This latter study also linked transcriptomic changes to the development of physiological function. However, considering the core role of rice as the major source of food for a large portion of the global population, and major global action to improve rice photosynthetic performance, the lack of information connecting early rice leaf development and transcriptomic data to metabolic and physiological changes represents a gap in our knowledge.

A complication arises in the study of leaf development in monocot grasses (such as rice) since investigations frequently take advantage of the basipetal nature of development in these organs (Nelson and Langdale, 1989, Li *et al*., 2010). Within grass leaves there is a gradient along the longitudinal axis of each leaf, with more differentiated cells towards the leaf tip while cells towards the base of the leaf remain in a proliferative state. This greatly facilitates experimental analysis since segments along the leaf comprise cells at different stages of differentiation, making it relatively straightforward to obtain tissue samples with different cell types at different stages of development. However, although there are many technical advantages to this approach, there is a conceptual challenge. The immature cells differentiating towards the proximal base of the leaf must experience signals from more distal, differentiated cells, in addition to signals reaching the leaf from other parts of the plant. Indeed, cell fate determination has already occurred once the cells have organised into files, as position is the primary determinant of cell fate in plants (Schiefelbein, 1994). This is distinct from the situation in a new leaf primordium on a meristem where *de novo* patterning must occur, with only very limited or no pre-pattern. Although some studies exist where this approach has been taken (Wang *et al*., 2013, Dechkromg, 2015, Miya *et al*., 2021), the data are more limited, reflecting the experimental challenges of primordium dissection and analysis.

*De novo* patterning in leaf development is most easily seen at the level of epidermal differentiation. Broadly, epidermal cells in leaves, including rice, can be split into pavement cells and more specialised cells. These specialised cells include guard cells and their associated subsidiary cells, and leaf hairs – also known as trichomes. Trichomes can be further characterised based on various features. While the terminology is not always consistent in the literature, rice displays two types of ‘stinging hair’ trichome (Maes and Goossens, 2010): ‘macrohairs’ (larger hairs seen only in the silica ladders, which are X-shaped structures regularly spaced between two walls, (Yamanaka S, 2009); and ‘prickle hairs’ (smaller, pointed trichomes which are more unevenly distributed over the epidermal surface, sometimes referred to in the literature as ‘microhairs’). Although these rice epidermal cell types have been well documented (Chaffey, 1983, Luo *et al*., 2012) exactly where and when these epidermal features arise during the *de novo* development of rice leaf primordia is yet to be precisely reported, although it is known that all epidermal features are present by the P4 stage of primordium development (Dechkromg, 2015). It is also unknown to what extent these epidermal patterns correlate with changes in internal leaf structure and function. With respect to function, as indicated above, leaf primordia are non-photosynthetic at initiation since they lack the biochemical and cell biological machinery to capture light and to utilise this energy and reducing power to fix CO_2_ into carbohydrate (van Campen *et al*., 2016). The process of acquiring photosynthetic capability is clearly a key step in leaf development and must be co-ordinated with a range of structural, physiological and biochemical changes at the level of the whole organ. In previous work, we addressed this issue in rice by performing a combined transcriptomic/chlorophyll fluorescence analysis of dissected primordia during very early stages of leaf development. These results allowed us to identify a phase in primordium development (P3/P4 transition) when a step change in physiology and structure occurred which accompanied the ability of leaf tissue to absorb and channel light energy towards photosynthesis (van Campen *et al*., 2016). This was co-ordinated with aspects of vascular and stomatal differentiation. However, although this work allowed us to identify a set of genes whose expression potentially underpinned the various structural and physiological processes observed, it did not provide direct information on the changes in biochemistry occurring. To address this issue, we have performed a metabolomic analysis of developing rice leaf primordia, focussing on the phase of development when the acquisition of photosynthetic capacity is established. To provide a higher resolution of developmental staging than previous work, we explored the use of a detailed developmental atlas of epidermal differentiation to see if there was a tight correlation of events on the leaf surface with internal changes in metabolism. This high-resolution developmental staging, combined with metabolomic analysis at the level of individual primordia, has allowed us to define a narrow developmental window during which a major re-organisation of leaf metabolism occurs.

## MATERIALS AND METHODS

### Plant Growth and Material

*Oryza sativa* ssp. *indica* var. IR64 was provided by IRRI. Seeds were germinated by submergence in water on filter paper in Petri dishes, then incubated in a SANYO growth cabinet, on a 12h 26°C / 12h 24°C light/dark cycle, PAR 2000μmol^-2^s^-1^ until the emergence of leaf 2, then transferred to a hydroponic system (van Campen *et al*., 2016). This was maintained in a Conviron BDR16 growth chamber. Conditions were maintained at 28°C and 60% humidity on a 12h/12h day/night cycle, with the light set at 700μmol^-2^s^-1^, (around 560μmol^-2^s^-1^ at seedling level). CO_2_ was maintained at ambient, which averaged at around 480ppm. The system consisted of a 6.5L opaque plastic container, filled with 5L of hydroponic growth media (1.4 mM NH_4_NO_3_, 0.6 mM NaH_2_PO_4_, 0.5 mM K_2_SO_4_, 0.8 mM MgSO_4_, 9 μM MnCl_2_, 1 μM (NH_4_)_6_Mo_7_O_24_, 37 μM H_3_BO_3_, 3 μM CuSO_4_, 0.75 μM ZnSO_4_, 70 μM Fe-EDTA). The level of media was maintained at 5L with water and replenished every 1-2 weeks. Seedlings were held in microfuge tubes with the bottoms removed, placed into polystyrene.

### Sample Staging, Dissection and Imaging

Leaf 5 primordia were staged non-invasively using the plastochron index based on length of leaf 3 (P1 – 4-20 mm, P2 - 25-70 mm, P3 – 75-110 mm, P4 120-140 mm, P5 – 150 mm or longer) (van Campen *et al*., 2016). Primordia were dissected with 25G and 30G hypodermic needles using a Leica MS5 dissection microscope and stored in water until imaging on the same day. Mature leaves were removed from the plant immediately prior to imaging.

Images were taken using a Hitachi TM3030 Plus Benchtop Scanning Electron Microscope. Images were captured in 15kV standard mode using secondary electron (SE) detection. Samples were blotted to remove excess water and then affixed to stubs using double-sided plastic tabs. All samples were imaged at −20°C using a cooling stage. Samples were scanned in ‘fast’ mode to find areas of interest and imaged in ‘slow’ mode.

All measurements were made using ImageJ Version 1.52a (Schindelin *et al*., 2015). Graphs were created using ggplot2 in R (R Core Team, 2019; RStudio Team, 2016; Wickham, 2016) (Versions: R – 3.6.2; RStudio – 1.1.453; ggplot2 – 3.2.1)).

### Mass Spectrometry

For metabolite extraction, staged, dissected primordia that had been flash frozen in liquid nitrogen, were freeze-dried for around six hours prior to extraction. Metabolites were extracted using the methanol-chloroform method based on a method by Overy et al. (Overy *et al*., 2005). For all extractions, LC-MS grade solvents (Honeywell) and distilled, deionised water were used. Samples were kept on ice and all centrifugation steps were refrigerated to 4°C.

Tissue samples of 1mg were ground in 10μL of an extraction medium comprising methanol/chloroform/water (2.5:1:1 by volume), left on ice for 5 minutes and then centrifuged at 14,000rpm. The supernatant was removed and the sample re-extracted by repeating the procedure with 5μL methanol/chloroform (1:1 by volume). To separate the supernatant into aqueous and chloroform phases, 3.5μL H_2_O and 2μL CHCl_3_ were added and the sample was then centrifuged at 14,000rpm for 15 minutes at 4°C, resulting in two distinct phases. The aqueous layer was removed and stored at −80°C and the chloroform phase was discarded. Small primordia where an accurate mass could not be determined were extracted as 1mg of tissue. Samples were stored at −80°C, and thawed on ice prior to use.

For ESI-TOF-MS, aqueous metabolite extracts were analysed using a Waters G2-Si mass spectrometer coupled to a Waters Acquity UPLC. The UPLC was used for automated injection of samples only. The injection volume of sample was 10μL and each sample was injected 3 times to obtain technical replicates. Spectral data were collected in negative ionisation mode, with a spectral window of 50 to 1200Da at a scan rate of 1 scan per second. A leucine enkephalin lock mass was run alongside the samples in order to check for drifts in mass measurement and accuracy over time. Peak lists were obtained as text files of accurate mass to 4 decimal places vs intensity. The three technical replicates per sample were combined using an in-house macro, as described (Overy *et al*., 2005). This procedure minimised noise in the samples and binned the data to 0.2 Da. All data was normalised to the total ion count. Multivariate analyses were performed using SIMCA (15.0.2) (Eriksson *et al*., 2006).

For MS/MS, from the multivariate analysis m/z ratios of interest were putatively identified using PubChem (Kim *et al*., 2019), with searches narrowed down by monoisotopic exact mass, and exclusion of compounds based on their lack of involvement in plant biochemistry. MS/MS analysis was run using direct infusion of the samples into a Waters G2-Si mass spectrometer. Using trap collision energy to fragment the ions of interest fragmentation patterns were obtained. These were compared to expected fragmentation patterns available online using the METLIN database (Guijas *et al*., 2018) or in the case where actual standards were available to real fragmentation patterns.

## RESULTS

### Creating an epidermal morphology atlas for early rice leaf development

With the aim of creating a system to allow a more precise developmental staging of rice leaf primordia prior to metabolomic analysis, we first focussed on analysing the patterning of the epidermis during leaf development. Mature rice leaves are characterised by a complex yet patterned series of elements on the epidermal surface (**Fig. 1A)**. These elements can be classified into macrohairs (**Fig. 1B**), pricklehairs (**Fig. 1C**) and stomatal complexes (**Fig. 1D**), in addition to the silica papillae visible in all these images. Preliminary investigations suggested that these different surface structures arose not only in a specific spatial pattern but also with a timing linked to the stage of leaf development. To characterise this further, we performed a detailed analysis of developing rice primordia utilising a previously described rice culture and staging system (van Campen et al., 2016). This system facilitates dissection of leaf primordia and grouping according to their plastochron age, as summarised in the introduction.

**Figure 1:**
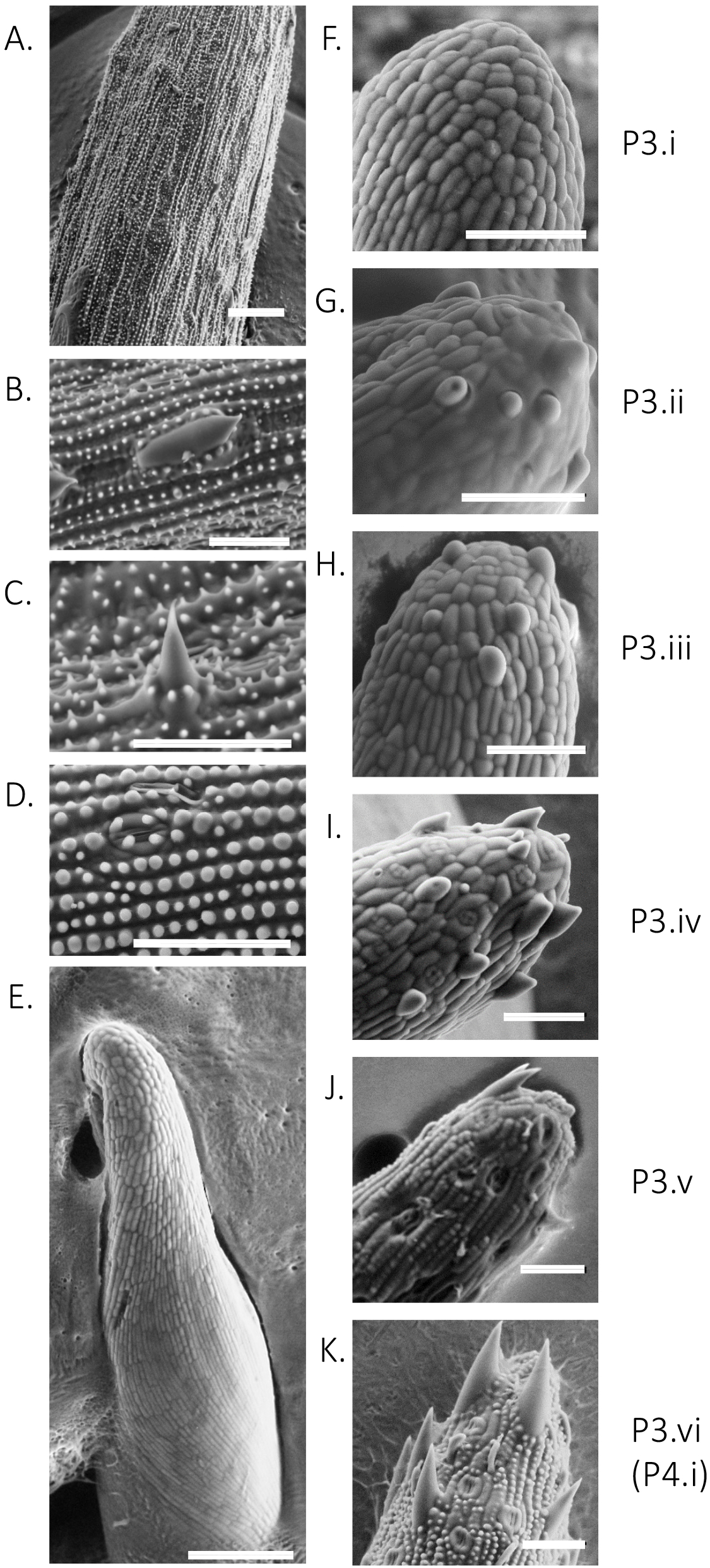
Patterning of epidermal features during early development of a rice leaf primordium. SEM images of **(A)** mature leaf epidermis, (**B)** macrohair, (**C)** pricklehair, (**D**) stomatal complex, (**E**) overview of a P3 primordium. Images of the tips of leaf primordia at stages (**F**) P3.i, (**G**) P3.ii, (**H**) P3.iii, (**I**) P3.iv, (**J**) P3.v, and (**K**) P3.vi. Scale bars: A and E, 1000µm; B-D, F-K 50µm.

Focussing on P3 primordia, our analysis revealed that at the beginning of this stage, although the epidermis was beginning to be patterned into the files of cells characteristic of mature grass leaves, there was no obvious differentiation of other epidermal structures (**Fig. 1E**), including at the leaf tip (**Fig. 1F**). This stage we defined as stage P3.i. The first epidermal structures appeared at the primodium tip and consisted solely of immature macrohairs (**Fig. 1G**, stage P3.ii). As the macrohairs matured, the precursor cell divisions presaging stomatal differentiation appeared at the primordium tip (**Fig. 1H**, stage P3.iii). Subsequent to this, prickle hairs started to form (**Fig. 1I**, stage P3.iv). During the next stage silica bumps (which later matured into papillae) appeared, by which time the macrohairs and stomata were fully differentiated and the prickle hairs were maturing (**Fig. 1J**, stage P3.v). By the end of P3 development, all surface elements of the mature leaf were fully visible (**Fig. 1K**, stage P3.vi) and the primordium tip epidermis was indistinguishable from that of the P4.i stage primordia. When the axial length of the primordia was plotted against sub-stage P3.i to P3.vi (as determined by analysis of the primordium tip), a close relationship was observed (**Fig. 2A**), indicating that primordium axis length can be used as a proxy for primordium tip differentiation. The data describing the developmental progression of tip epidermal differentiation in P3 stage primordia are summarised in schematic form in **Fig. 2B**.

**Figure 2:**
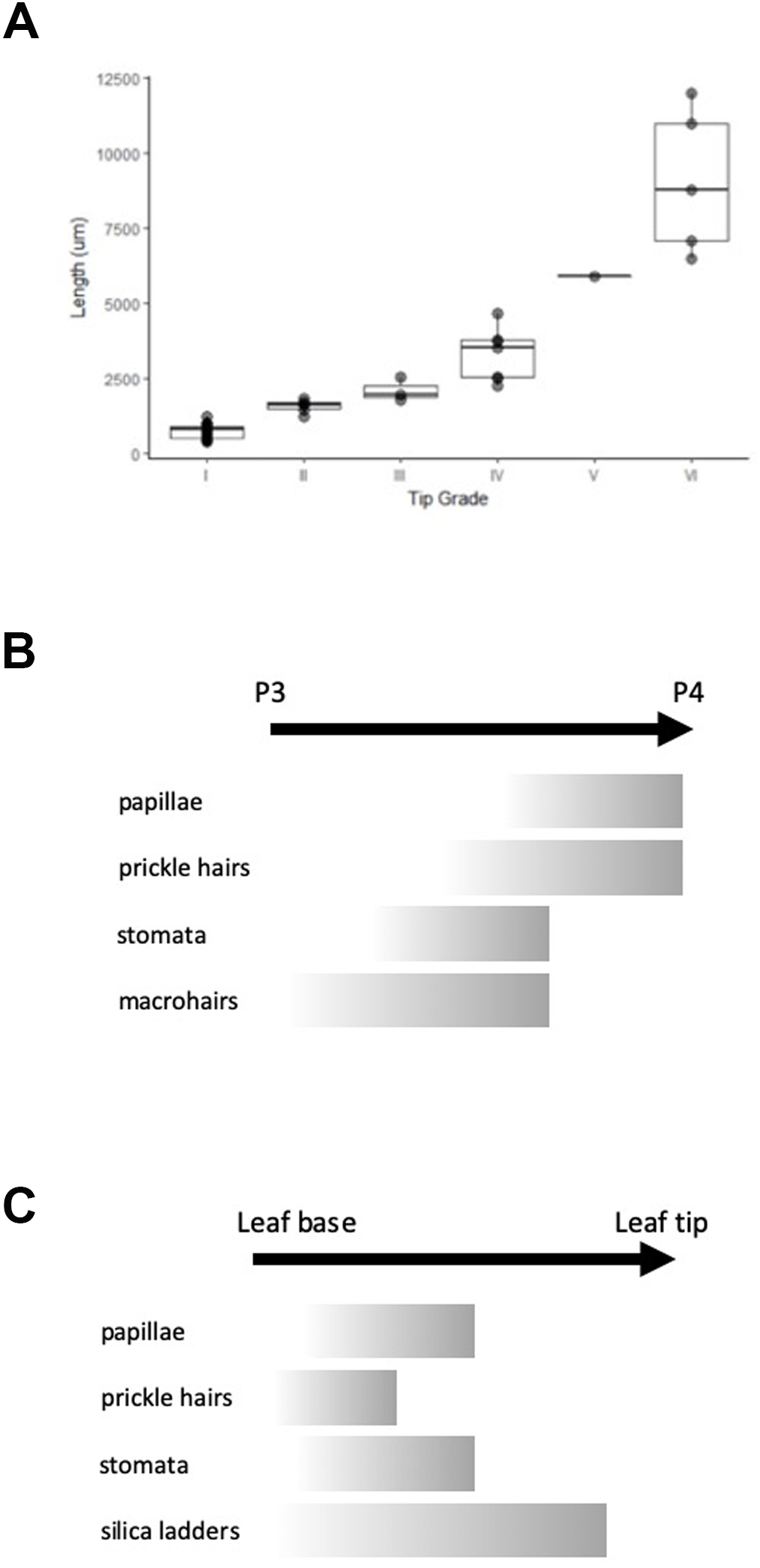
Epidermal patterning at the primordium tip correlates with growth. **(A)** Axial length of leaf primordia against primordium tip developmental stage. Boxplots show the first quartile, mean and third quartile for each stage. n = 23. **(B)** Schematic summarising the spatial/temporal appearance of different epidermal features during the development of leaf primordia from P3 to P4 stage. **(C)** Schematic summarising the spatial/temporal appearance of different epidermal features along the axis of a mature leaf from base to tip.

### Analysis of primordia reveals a rapid major shift in metabolite profile during the late P3 stage of development

In initial experiments to investigate any change in metabolic profile during early leaf development, primordia were dissected and staged into P3, P4 and P5 groups (van Campen *et al*., 2016). The primordia were extracted into solvent and the polar phase was analysed by direct infusion electrospray ionisation-time of flight-mass spectrometry to produce a metabolite fingerprint which depicts a snapshot of the metabolome at each stage of development. Multivariate analysis via principal component analysis (PCA) indicated that bulked P3, P4 and P5 primordia could be distinguished from each other based on metabolite profile **(Fig. 3A**). When the same analysis was performed using the metabolite profiles of individual dissected primordia which fell into the three main plastochron stages, and analysed using PCA, a similar picture emerged (**Fig. 3B**), indicating that the approach was sufficiently sensitive to distinguish extracts from single primordia. When the analysis was then extended to include individual, dissected primordia staged according to the system described in Fig. 1 and Fig. 2 (i.e., a higher developmental resolution), a more complicated picture emerged (**Fig. 3C**) with samples appearing intermediate between the P3, P4 and P5 stages. Nevertheless the sub-groups of primordia tended to be distinguishable from each other, suggesting that the intermediate primordia had metabolite profiles which lay between the main stages identified in Figs. 3A,B, thus potentially reflecting a transition in metabolism.

**Figure 3:**
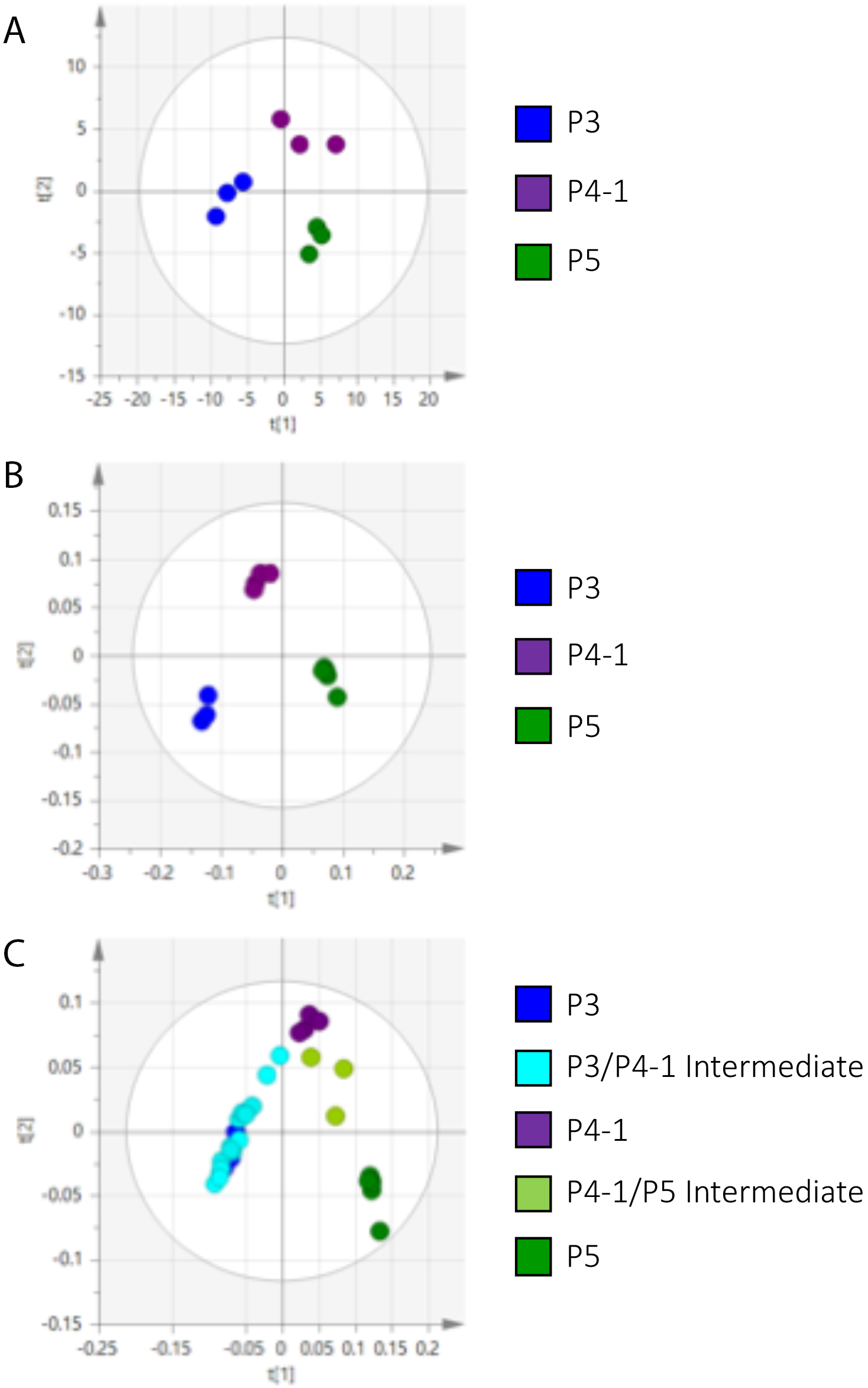
PCA of metabolite profiles for bulked and individual primordia allows discrimination of developmental stages. **(A)** PCA plot of samples comprising bulked primordia at P3, P4.i or P5 stage, as indicated. **(B)** PCA plot of samples from individual primordia falling into stages P3, P4.i or P5 stage, as indicated. **(C)** PCA plot of samples from individual primordia staged from P3.i through to P5 using the epidermal tip index described in Figure 1.

To make an initial identification of which metabolites might be allowing the separation of the developmental stages described in Fig. 3, we performed pairwise unsupervised PCA analysis of P3-P4 (**Fig. 4A**) and P4-P5 staged primordia (**Fig. 4B**), and interrogated the associated loading plots (**Fig. 4C, D**). When the lead loading plot metabolites were ranked, metabolites associated with the tricarboxylic acid (TCA) cycle were noticeably present in both the P3-P4 (**Fig. 4E**) and P4-P5 comparisons (**Fig. 4F**).

**Figure 4:**
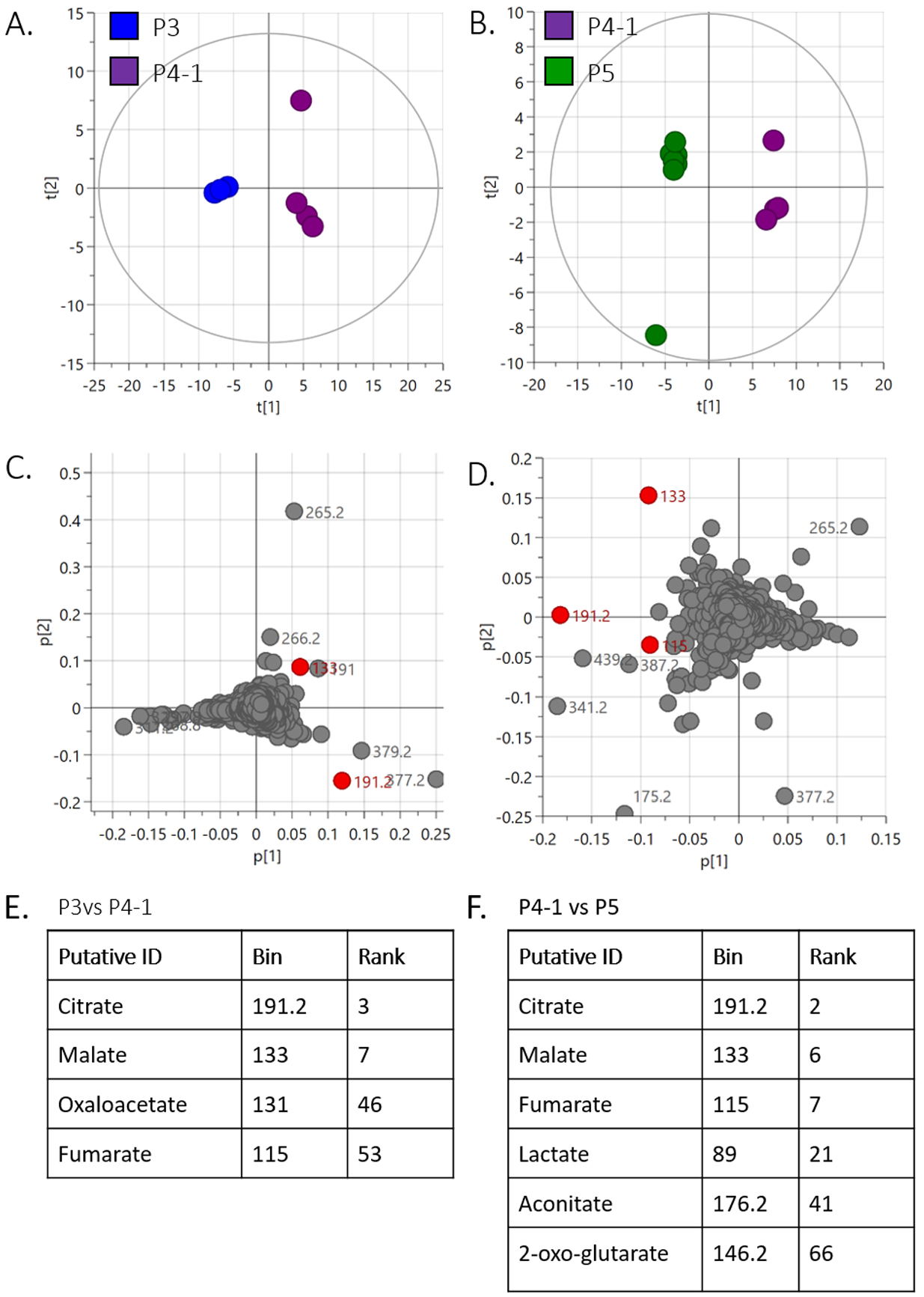
Pairwise comparisons of metabolite profiles during primordium development identifies TCA metabolites as discriminating ions. **(A, B)** PCA plots of pooled primordia samples at P3 and P4.i stages **(A)** and P4.i and P5 stages **(B)**, as indicated. **(C**,**D)** Loadings plots for the respective PCA plots shown in (A,B). M/z ions indicated in red indicate metabolites having a major influence in distinguishing samples by this approach and whose putative identity is linked to the TCA cycle. **(E**,**F)** Ranking of TCA metabolites for **(E)** the P3-P4.i analysis shown in (C), and **(F)** for the P4.i-P5 analysis shown in (D).

### Evidence for an altered TCA pathway configuration during the late P3 stage of leaf development

To investigate this observation further, we identified all the m/z ions related to TCA metabolites and quantified their level relative to total ion count using the primordium staging described in Fig. 1 and Fig. 2. Strikingly, at the transition from P3.iv to P4.i stage there was a dramatic increase in the relative levels of citrate, fumarate and malate (**Fig. 5A, D, E**). In contrast, the relative level of ketoglutarate fell during this transition (**Fig. 5B**), whereas oxaloacetate level rose to a peak at P3.iii/P3.iv before falling during the P4 and P5 stages (**Fig. 5F**). Levels of succinate were variable (**Fig. 5C**), with if anything a slight decline from early P3 to later P3/P4 stage. To confirm that the m/z ions identified in our metabolomic analysis did indeed represent TCA metabolite, a series of MS/MS analyses were performed which substantiated the identity of the m/z ions as TCA metabolites (**Supp. Table 1**). The relative changes in TCA metabolite level during primordium development are summarised in schematic form in **Fig. 5G**, with the major changes in metabolite level most apparent when comparing the P3.iv and P4.i stages of primordium development.

**Figure 5:**
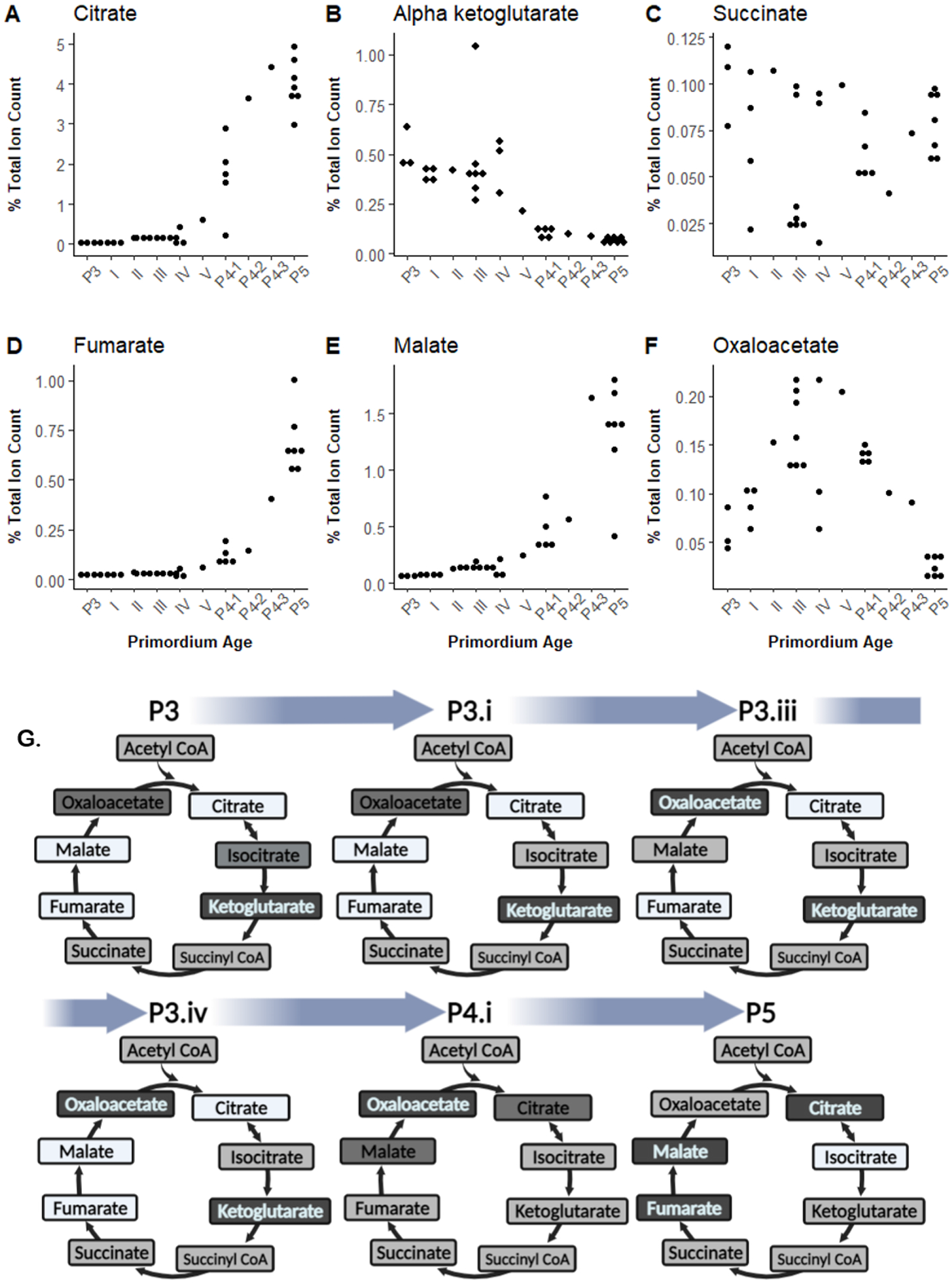
A non-uniform shift in TCA metabolism occurs during the late P3 stage of leaf development. **(A-F)** Relative ion counts detected in individual primordia at different developmental stages for (A) citrate (B) alpha ketoglutarate (C) succinate (D) fumarate (E) malate (F) oxaloacetate. **(G)** Schematic of the relative levels of TCA metabolites at different stages of primordium development. Darker shading indicates relatively high ion counts, lighter shading indicates lower relative ion counts.

### Comparison of metabolite and transcript levels during early leaf development supports a presaging of altered metabolism by altered gene expression

To investigate the extent to which the changes in metabolite profiles described above were presaged at the transcriptional level, we interrogated an RNAseq database covering the P3, P4 and P5 stages of rice leaf development (van Campen et al., 2016). **Fig. 6** shows a summary of the overall changes in metabolite and mRNA levels according to different expression profiles. In this scheme, transcript and metabolite changes have been classified based on their pattern during P3-P4-P5 primordium development, as indicated. For example, “UP1” and “DOWN1” collates changes in level that occurred predominantly during the P3 to P4 transition, while “UP2 and DOWN2 identifies groups of metabolites or transcripts where changes in level predominantly occurred on the transition of P4 to P5.

**Figure 6:**
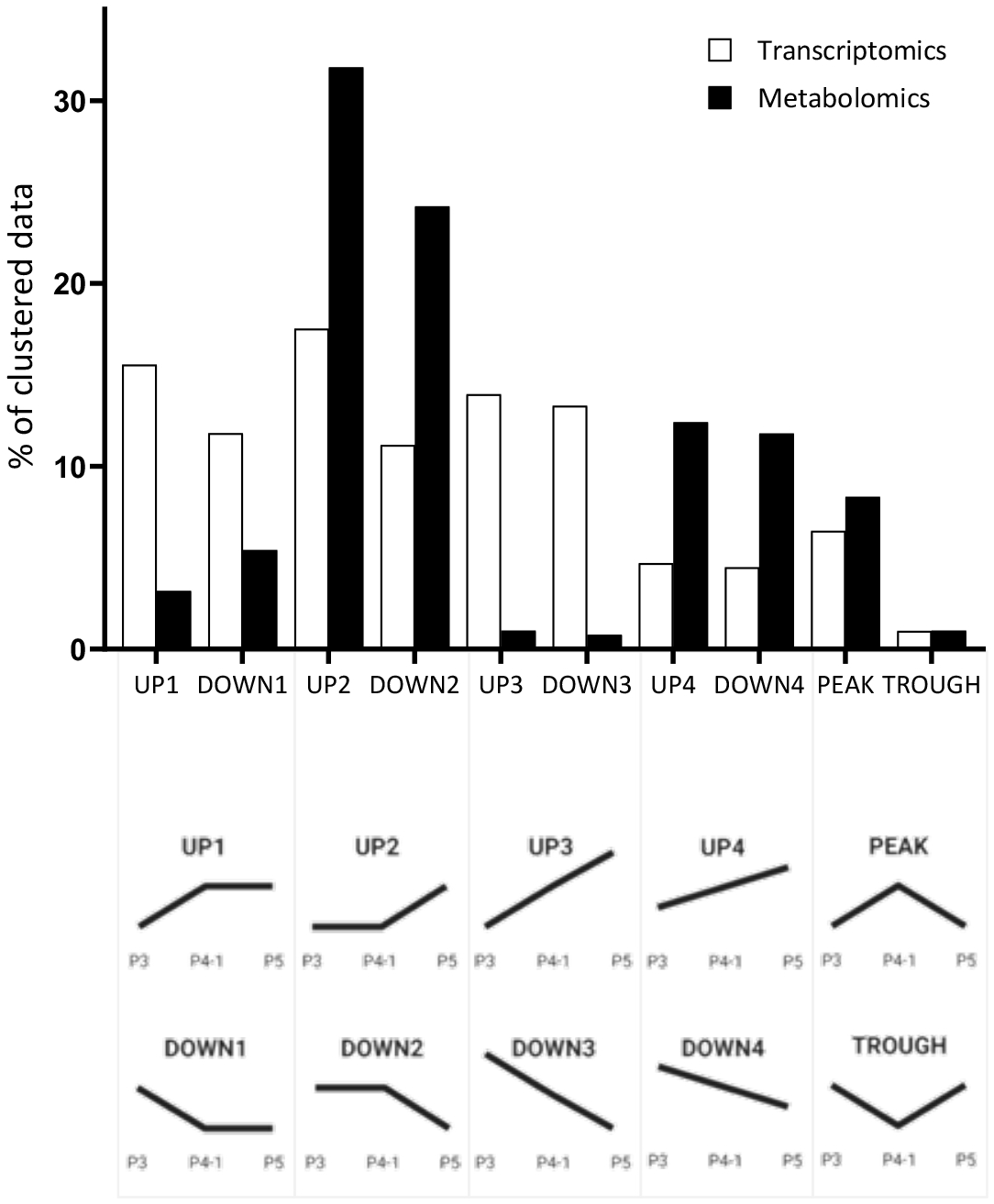
Shifts in transcription at the P3-P4 transition presage shifts in metabolism at the P4-P5 transition in primordium development. Relative changes in cluster pattern for transcriptome data (white bars) and metabolome data (black bars) over the transition from P3 through to P5 stages of development. The patterns have been defined as indicated in the schematics below the graph, adapted from van Campen et al (2016), with UP1 and DOWN1 indicating major relative change at the P3 to P4 transition, whereas UP2 and DOWN2 indicate major relative change at the P4 to P5 transition. UP3/DOWN3 indicates relatively rapid, continuous change from P3 to P5, whereas UP4/DOWN4 indicate more gradual yet continuous change over the P3 to P5 transition. PEAK and TROUGH indicate patterns where data cluster either had a maximum or minimum level at P3 stage.

This analysis indicated that although some changes in metabolite profile were clearly happening during the P3 to P4 transition (including, for example, the changes in TCA metabolites reported above) the bulk of metabolite changes occurred later in leaf development (P4 to P5 transition; UP2, DOWN2, UP4, DOWN4 classifications). In contrast, changes in transcript level (either up or down) were more evenly distributed throughout the developmental stages analysed, with relatively large changes compared to the recorded metabolite changes occurring during the P3 to P4 transition (UP1, DOWN1 classifications). This fits to a scenario where transcript changes broadly presaged changes in metabolism and where the anatomical, biochemical and physiological events initiated towards the very end of the P3 stage set in train the more major changes in metabolism recorded during P4 and P5 stages of development as the leaf approached maturity.

## DISCUSSION

Our previous data identified the P3/P4 transition in early rice leaf development as the stage when major changes in rice physiology were occurring, notably the acquisition of the capability to capture light energy and channel it into the pathway required to drive the capture of CO_2_ via photosynthesis (van Campen et al., 2016). The data presented here extend this analysis to show that the previously documented changes during early leaf primordium development at the level of structure, gene expression and physiology are co-ordinated with changes in metabolism. In particular, the creation and implementation of a primordium tip differentiation atlas, along with the use of metabolomic analyses of single primordia, allowed us to identify a relatively narrow phase in primordium development at the end of the P3 stage when major reconfiguration of primary metabolism occurs. Taken in the context of previous data, these results indicate a relatively rapid switching of core metabolism as primordia gain autotrophic capability.

### Use of epidermal features to stage early leaf development

The majority of studies of leaf development in rice and other monocot crops have considered the classical spatial developmental trajectory in which tissue towards the distal tip of a mature leaf is fully differentiated while tissue towards the proximal base of a leaf is less differentiated. While a perfectly valid approach, from a developmental point of view it has two limitations. Firstly, the spatial resolution used in the analysis is often performed at the mm to 10’s mm range whereas developmental paracrine signals generally act over the range of 10’s – 100’s µm. Secondly, differentiation at the proximal base of maturing leaves will inevitably be informed by the mature distal tissue, which will (by definition) have a complex pre-pattern and, moreover, be physiologically active, generating a wealth of potential information signals which may influence the patterning/differentiation process in the “naïve” tissue at the base. In contrast, the approach taken here examined the *de novo* leaf primordium which will have very limited pre-pattern and very limited information flow from differentiated cells in the rest of the leaf. It is thus closer to addressing the fundamental question of how a naïve volume of tissue develops the complex yet integrated structural, biochemical and physiological framework required for the formation of a functional leaf (Nelissen *et al*., 2016). The order of patterning of epidermal elements described here is a case in point. The spatial/temporal order observed in developing primordia (**Fig. 2B**) is similar but distinct from that observed along the proximal-distal axis of mature rice leaves (**Fig. 2C)**. This supports the proposal that although the signalling processes involved are probably highly analogous, some elements of *de novo* pattern generation in developing leaf primordia are distinct from those observed in systems where some form of pre-pattern exists.

The functional significance of the patterning of the majority of the epidermal surface structures used to stage the leaf primordia here is unclear. Although roles in, e.g., creating a surface microenvironment to aid photosynthesis/restrict water loss and/or roles in anti-herbivory or inhibition of pathogen attack are well-established (Glover, 2000), why the structures appear in the precise spatial/temporal pattern reported here is more obscure. Identification and functional characterisation of mutants in which these patterns are disrupted might shed light on this issue. Irrespective of the function of the patterns, the ability to visually and class dissected primordia in a robust staging system enabled us to demonstrate that distinct developmental stages of early leaf development were characterised by consistent patterns of metabolism.

### Analysis at single primordium resolution reveals a major reconfiguration of metabolism during a narrow phase of early leaf development

Metabolite fingerprinting is a powerful tool that has proven useful in a wide area of biology, including medical diagnostics (Wei *et al*., 2021), discriminating plant populations (Basile *et al*., 2018; Xiao *et al*., 2018) and tracking plant pathogen infection (Rubert *et al*., 2017) as just a few examples. In the field of developmental biology, metabolite fingerprinting has proven a useful tool to differentiate between different embryonic stages in both *Drosophila melanogaster* and zebrafish (An *et al*., 2014, Dhillon *et al*., 2019). The sensitivity and resolution of mass spectrometry techniques continues to improve so that it is now possible to get reliable data from very limited amounts of tissue. Here we report on metabolomic analysis at the resolution of single primordia, which we believe is a novel application of the technique. The tight clustering of data points according to primordia at distinct stages along the developmental trajectory indicates the robustness of the approach, suggesting that it will be readily applicable to other plant systems. Clearly no single metabolomic approach can detect all metabolites in a complex biological sample. Of particular relevance to this study, due to the issues of volatility and the presence of phosphate groups, many Calvin-Benson cycle metabolites are unlikely to be efficiently detected with the global approach taken here. Targeted metabolomics of the Calvin-Benson cycle would require significantly more pre-processing, such as linking to LC-MS or isotope labelling (Arrivault *et al*., 2009). We thus view the data here as an initial validation of the metabolomics approach being able to distinguish primordia during a developmental trajectory, with the results identifying a window during which major changes in metabolism occur. Our approach has allowed some components underpinning this change to be identified (described below) but should not be seen as identifying all the changes that are occurring, which will require a more targetted approach.

The TCA cycle is core to eukaryotic primary metabolism, providing energy, reducing power and carbon skeletons for a very wide range of metabolic processes. In plants, some of the TCA organic acids also function as osmolytes, accumulating to relatively high levels in the vacuole to help drive growth, as well as shuttling between cells to co-ordinate metabolism (Sweetlove *et al*., 2010, Geigenberger and Fernie, 2014, Igamberdiev and Eprintsev, 2016). A dramatic increase in the accumulation of some TCA metabolites occurred during a narrow phase of primordium development (P3iv to P4.i) (e.g., malate, citrate, fumarate) whereas others declined sharply (e.g., ketoglutarate). The functional significance of these shifts awaits elucidation, but they are consistent with the hypothesis that a major general change in metabolism is occurring at this stage. Since our previous work identified the same stage undergoing a transition in physiology from heterotrophy to autotrophy (van Campen *et al*., 2016), the results substantiate the hypothesis that the late P3 primordium stage of development is a key stage in rice leaf development. To give some sense of scale, this transition occurs while primordium length increases from approximately 5 to 7.5mm.

This relatively rapid transition contrasts with the conclusion by some other studies which have exploited the developmental gradient along monocot grass leaves to investigate the changes in metabolism occurring as leaf differentiation occurs (Pick *et al*., 2011, Wang *et al*., 2014), with more gradual gradients of metabolic change being observed. We suggest this probably simply reflects the developmental differences in the two systems, along with the spatial resolution analysed (the most proximal tissue in the leaf-segmentation approach already tending to be slightly green, as opposed to the dissected early P3 primordia reported here which are non-photosynthetic). It is noticeable that an analysis of Arabidopsis leaf development also suggested a relatively rapid acquisition of photosynthetic differentiation linked to a co-ordinated exit from cell cycle (Andriankaja *et al*., 2012). The division status of the cells in the rice primordia described here was not analysed, but would be an interesting avenue of future investigation.

Finally, the results described here (along with others in the literature) indicate that although important aspects of rice leaf structure and function are laid down at a relatively early growth stage (axial length circa. 5mm), there is a phase before this when, presumably, major elements of structure/function formation are very plastic (since they have not yet been formed). Understanding more about this phase of developmental plasticity may be advantageous for efforts aimed at engineering rice leaf structure and function (Ort *et al*., 2015, Ermakova *et al*., 2020).

## ACKNOWLEDGEMENTS

This work was supported by an iCASE award to NC/AJF as part of the BBSRC White Rose DTP (BB/M011151/1), working with the International Rice Research Institute (IRRI).

## AUTHOR CONTRIBUTION

Investigations and data analysis were performed by NC, JW and JP. Project was conceived by AF and PQ, with supervision by AF, LS and HW (Sheffield) and PQ (IRRI). The original draft was by AF, NC and LS, with all authors contributing to review, editing and visualisation.

## DATA AVAILABILITY

The data supporting the findings in this study are available from the corresponding author (Andrew Fleming), upon request.

## FIGURE LEGENDS

**Supplementary Table 1: MS-MS data confirms the identity of TCA metabolites**

